# Detection of Blood Volume Reduction and Vasoconstriction Following Focused Ultrasound Blood-Brain Barrier Opening Using Ultrasound Flow Imaging

**DOI:** 10.1101/2023.10.19.563176

**Authors:** Sua Bae, Stephen A. Lee, Elisa E. Konofagou

## Abstract

Microbubble-mediated focused ultrasound (FUS) offers a non-invasive treatment for transient and localized blood-brain barrier (BBB) opening for drug delivery or immunostimulation. It is known that FUS-induced BBB opening is accompanied by blood flow changes, vasoconstriction, and vasodilation, as validated by optical microscopy through a cranial window. In this study, we introduce a novel method for quantifying vascular changes after FUS-induced BBB opening by employing ultrasound flow imaging in mice. We acquired pre-FUS and post-FUS ultrasound flow images with the same microbubble concentration in the brain. Contrast-enhanced power Doppler (CEPD) images and ultrasound localization microscopy images were obtained to evaluate changes in cerebral blood volume and vessel diameter at the sonicated region of the brain. Our findings demonstrate that FUS leads to a reduction in blood volume at the treated region, with vasoconstriction being more dominant than vasodilation. Furthermore, we show that transcranial CEPD can detect local blood reduction following FUS, which spatially coincides with the edema region identified in T2-weighted MRI. Our findings suggest that ultrasound flow imaging has the potential to serve as a cost-effective and immediate monitoring tool for evaluating the safety and efficacy of FUS-induced BBB opening.

## Introduction

Microbubble-mediated focused ultrasound (Mb-FUS) is a promising non-invasive treatment for the transient and localized blood-brain barrier opening (BBBO) to enhance drug delivery (Konofagou, 2012; McMahon et al., 2021) and promote immune responses (Ji et al., 2021; Kobus et al., 2016; Leinenga & Götz, 2015). In this treatment, microbubbles are administered systematically, and focused ultrasound (FUS) induces rapid and nonlinear oscillations of microbubbles within a targeted volume of the brain. These oscillations, known as cavitation, apply mechanical forces to the blood vessel walls and capillaries, causing the transient disruption of tight junctions between endothelial cells and the increase of transcytosis and fenestration (Roovers et al., 2019; Sheikov et al., 2008; Karakatsani et al. 2023).

Some studies using optical microscopy showed that Mb-FUS induces transient vessel constriction and dilation in the brain through the cranial window (Cho et al., 2011; Raymond et al., 2007). Cho et al. found that vasoconstriction is more prevalent than vasodilation in mice and the constrictions were maintained for 5–15 min. A. Burgess et al. showed that leakage of the dye through the vessel walls was accompanied by vasodilation which was occasionally preceded by rapid vasoconstriction in transgenic mice (Burgess et al., 2014). Although optical imaging can visualize dye leakage and changes in vessel diameter, it has limited penetration depth, making it unsuitable for transcranial monitoring without skull removal.

On the other hand, ultrasound imaging has a higher penetration depth and Doppler imaging has been used for transcranial blood flow imaging for cerebrovascular structure and function (Willie et al., 2011). Furthermore, microbubbles can be used as a contrast agent to enhance imaging sensitivity through the skull (Errico et al., 2016). Ultrasound localization microscopy (ULM) with microbubbles can provide high-resolution microvascular imaging below the ultrasound diffraction limit (Errico et al., 2015). This is achieved by localizing the bubbles from the hundreds of thousands of low-resolution images. Despite these advantages of ultrasound flow imaging, to the best of our knowledge, Doppler imaging or ULM has not yet been utilized for BBB opening monitoring or assessment.

Currently, MRI is the most commonly used method to confirm the safety and efficacy of FUS-induced BBB opening in vivo without the need for sacrificing. In both preclinical and clinical studies, the animals or subjects are transferred to the MRI to obtain various scans including T2-weighted, T2*-weighted, FLAIR, or SWI images for detecting edema or hemorrhage, and contrast-enhanced T1-weighted images for detecting the BBB opening (Chen et al., 2021; Mainprize et al., 2019; Pouliopoulos et al., 2021). However, these scans are not feasible to obtain simultaneously with Mb-FUS or immediately after Mb-FUS due to the time required for setting up the MRI scan, unless the FUS setup was equipped with the MR scanner.

For real-time monitoring of FUS treatment, cavitation dose detection or mapping is generally used (Pouliopoulos et al., 2021, Bae et al., 2023). Cavitation mapping visualizes the location and intensity of cavitation activity at the targeted area during the sonication (Coviello et al., 2015), allowing for assessing the effectiveness of the FUS and optimizing the acoustic parameters. While many preclinical and clinical studies have shown a high correlation between cavitation dose and the size of opening or damage (Bae et al., 2023; Jones et al., 2020; Yang et al., 2019), cavitation monitoring only measures the acoustic energy generated by microbubbles rather than directly showing their biological effects. Given the varied responses of different tissue and vessel types to microbubble cavitation, blood flow imaging would offer more direct information about the resulting bioeffects.

In this paper, we propose ultrasound flow imaging as a monitoring tool for Mb-FUS in BBB opening treatment. We establish a method to quantify the vascular changes before and after BBB opening.

## Methods

### Experimental setup

We employed two different experimental setups for open-skull and transcranial experiments. In the open-skull experiments, imaging and therapy were performed using the same linear array transducer (L22-14vXLF; number of elements: 128, center frequency: 15.6 MHz) as illustrated in Figure 1A. For BBBO with the imaging transducer, we used the theranostic ultrasound sequence as described in (A. Batts et al., 2022), utilizing electronically-focused ultrasound with a short pulse. Given that the transmit frequency of the probe we used here was 10 times higher than our previous theranostic ultrasound frequency (15.6 MHz vs. 1.5 MHz), the focal size was only ∼0.1 mm in width with an F-number of 1. To compensate for the small focal size, we transmitted 5 foci spanning 0.5 mm in the lateral direction. The sonication sequence and parameters are presented in Figure 1A and Table 1. The number of bursts was 60, and the burst repetition frequency was 0.5 Hz (i.e., 2 min of total sonication time). In each burst, 100 pulses per focus were sonicated with a pulse repetition frequency of 1 kHz. The 5 pulses for the 5 foci were transmitted with a between-foci interval of 17 μs considering the round-trip time for imaging depth of 10 mm. The mechanical index of the focused beam was 0.6, and the pressure was 2.3 MPa. The left hippocampus and thalamus were targeted for BBBO at 2 mm caudal from bregma. A research ultrasound system (Vantage 256, Verasonics Inc., Kirkland, WA, USA) was used for controlling the transmit sequence and acquiring the data.

**Table 1.** Parameters for FUS sonication for BBBO in the open-skull and transcranial experiments.

|  | Open-skull experiment | Transcranial experiment |
| --- | --- | --- |
| Transducer | Linear array probe (L22-14vX-LF) | Single-element, spherical transducer |
| Frequency | 15.6 MHz | 1.5 MHz |
| Focal depth | 5 mm | 60 mm |
| F# | 1 | 1 |
| Pressure | 2.3 MPa | 150-450 kPa (derated) |
| Assumed skull-induced attenuation | N/A | 20% |

**Figure 1:**
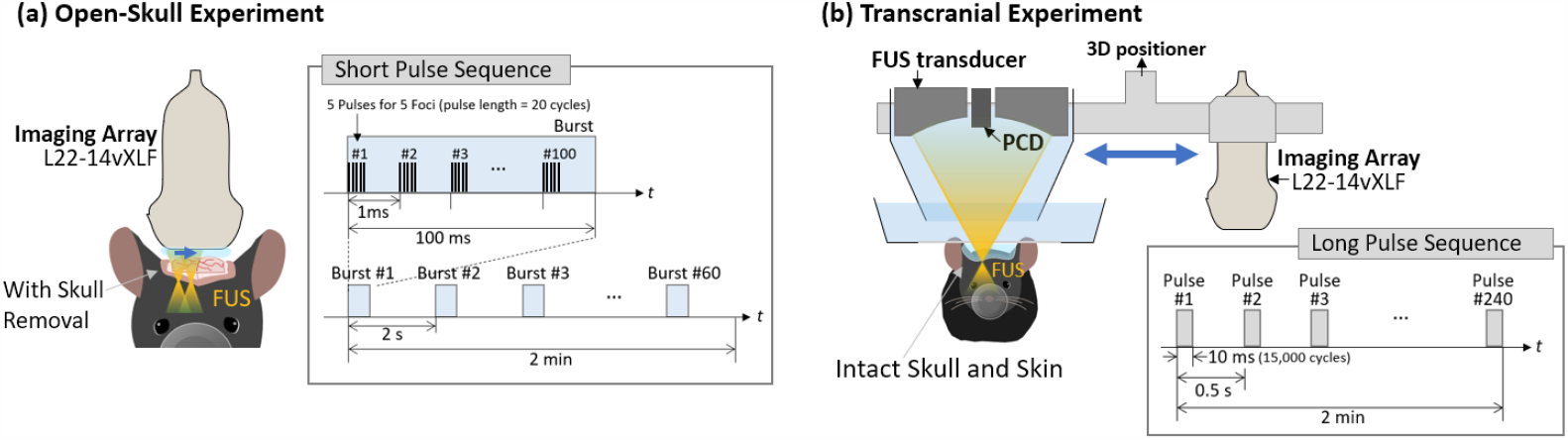
Experimental setups and pulse sequences for (a) open-skull experiment and (b) transcranial experiment.

In transcranial experiments with intact skin and skull, a conventional spherical FUS transducer (diameter: 60mm, focal depth: 60 mm) was used for a long pulse sequence for BBBO and the same 15.6-MHz linear array transducer was used for transcranial imaging (Figure 1B). The array was placed on the mouse head for pre-FUS imaging and then was replaced with the FUS transducer for BBBO using the 3D positioner. Immediately after FUS, the array was placed back in the same position for post-FUS imaging. For therapeutic sonication, a 10-ms long pulse was sonicated for 2 min with an interval of a half-second (Figure 1B). The FUS frequency was 1.5 MHz, and the derated pressure of FUS ranged from 150 to 450 kPa, assuming skull-induced attenuation of 20%.

### Acquisition and Reconstruction of Flow Images

In both the open-skull and transcranial experiments, we used the same imaging sequence to acquire flow images before and after Mb-FUS (Figure 2). Ultrasound flow images were obtained after a bolus injection of microbubbles and a low-resolution CEPD (pixel size: 0.2mm×0.2mm) was displayed for real-time monitoring of the bubble concentration in the mouse brain. With another bolus injection, FUS was sonicated for 2 minutes to open the barrier at the left hippocampus and thalamus. Power cavitation imaging was also performed by the array transducer during FUS in the open-skull experiment as in the previous studies (A. Batts et al., 2022; A. J. Batts et al., 2023; M. T. Burgess et al., 2018). In the transcranial experiments, the cavitation dose was monitored by using the passive acoustic detector (PCD) shown in Figure 1B. After sonication, post-FUS flow images were obtained by administering additional microbubbles until the CEPD intensity matched that of the pre-FUS images.

**Figure 2:**
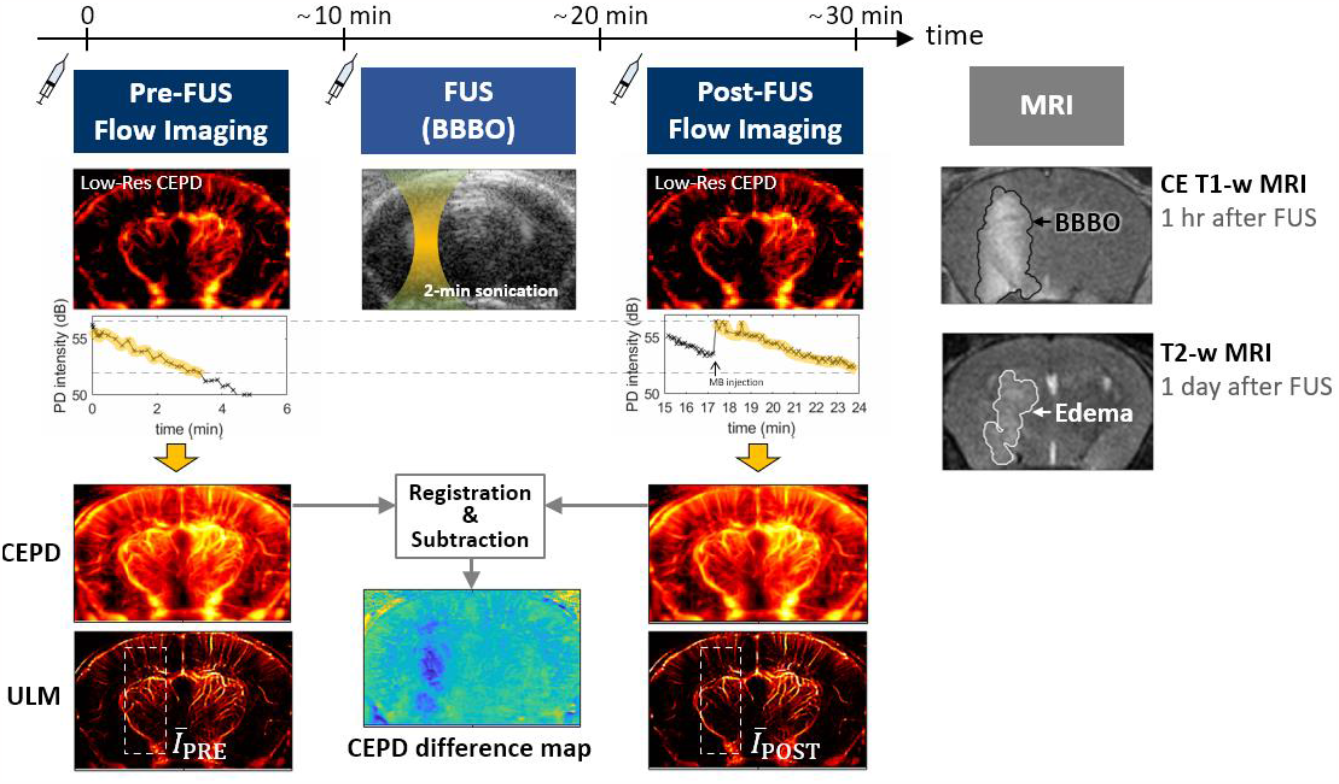
Acquisition and process of contrast-enhanced power Doppler (CEPD) and ultrasound localization microscopy (ULM) imaging before and after Mb-FUS.

For flow imaging, plane wave compounding with 9 steering angles was employed to obtain a set of 500 frames with an effective frame rate of 500 kHz (Table 2). Multiple sets of data were obtained for 5–10 min. As the flow signal intensity is highly dependent on the microbubble concentration in the brain, we used the data sets within the same range of CEPD intensity for reconstructing pre-FUS and post-FUS images as indicated by the yellow highlight in Figure 2.

**Table 2.**
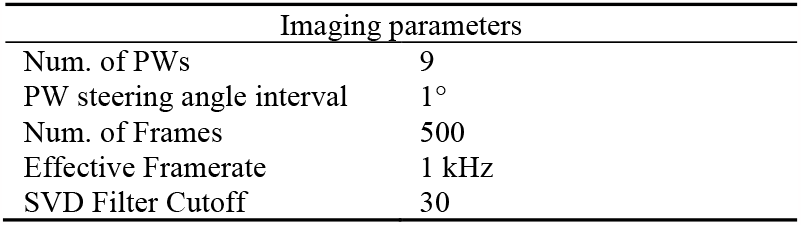
Parameters for flow imaging in both open-skull and transcranial experiments.

Inphase-quadrature (IQ) beamformer was used to form the ultrasound image (Bae et al., 2021), and singular value decomposition (SVD) filtering was applied to the IQ-beamformed images removing the tissue and breathing motion (Errico et al., 2016). High-resolution CEPD (pixel size: 50µm×50µm) and ULM (pixel size: 6.25µm×6.25µm) images were reconstructed offline. We obtained CEPD images by squaring the pixel intensity of the filtered images and averaging all the frames of multiple data sets. In the case of ULM, IQ beamformed images were reconstructed with a pixel size of 25µm×25µm. They were then processed by SVD filtering, interpolated by a factor of 2, and deconvoluted using a Gaussian filter (standard deviation: 50µm×50µm). To localize microbubbles, the *regionalmax* function in MATLAB was employed after thresholding at the 0.8 quantiles of pixel intensity and interpolating by a factor of 2. The final ULM image was obtained by averaging all the localized images of multiple data sets.

### Analysis of Flow Images

Flow intensity was measured by averaging the intensity within a region-of-interest (ROI) centered at the FUS focus. Then, the averaged intensity was normalized by that of the contralateral region. The blood volume change after sonication was measured by subtracting the normalized intensity of pre-FUS image from the normalized intensity of post-FUS image. The vessel segments that were well-reconstructed in both pre-FUS and post-FUS images and did not overlap with other vessels were selected for the vessel diameter measurement. Fifteen cross-section profiles perpendicular to the vessel direction were obtained along the length of 30 μm with an interval of 2 μm. The diameter of each segment was estimated by averaging the cross-section profiles and measuring its full-width half-maximum.

### MRI

MR scan was performed for each mouse to confirm BBB opening and assess the edema (9.4T Ascend, Bruker Medical, Billerica, MA). Contrast-enhanced T1-weighted MRI was obtained approximately 1 hour after Mb-FUS and 30 min after the intraperitoneal injection of a gadolinium-based MR contrast agent (Omniscan, Princeton NJ; 0.2 mL per mouse). Contrast-enhanced T1-weighted images were used for the detection and quantification of BBB opening. T2-weighted images were also obtained 1 day after Mb-FUS without contrast enhancement for assessment of edema.

### Animals

The animal studies were conducted in compliance with the guidelines established by the Institutional Animal Care and Use Committee (IACUC) of Columbia University and were approved by the same committee. Wild-type 6–10 week-old mice (C57BL/6, male; The Jackson Laboratory, Bar Harbor, ME, USA) were used in the study. For the open-skull experiments, a total of four mice were employed, while ten mice were used for the transcranial experiment. The latter group of ten mice was further divided into four subgroups, each exposed to different acoustic pressures: N = 2 (150 kPa), N = 3 (250 kPa), N = 3 (350 kPa), and N = 2 (450 kPa).

## Results

### Flow intensity reduction and diameter after Mb-FUS in the open-skull experiment

Figure 3A presents the power cavitation image during the sonication for BBBO through the cranial window. The power cavitation image showed the microbubble activities at the sonicated region. BBBO for each mouse was confirmed by T1-weighted MRI (Figure 3B). ULM images at the treated brain region before and after FUS are presented in Figure 3C, indicating a decrease in signal intensity after the sonication (white arrow heads). The normalized flow intensity within the windowed area decreased after Mb-FUS in all four mice with an average of -10% (Figure 3D).

**Figure 3:**
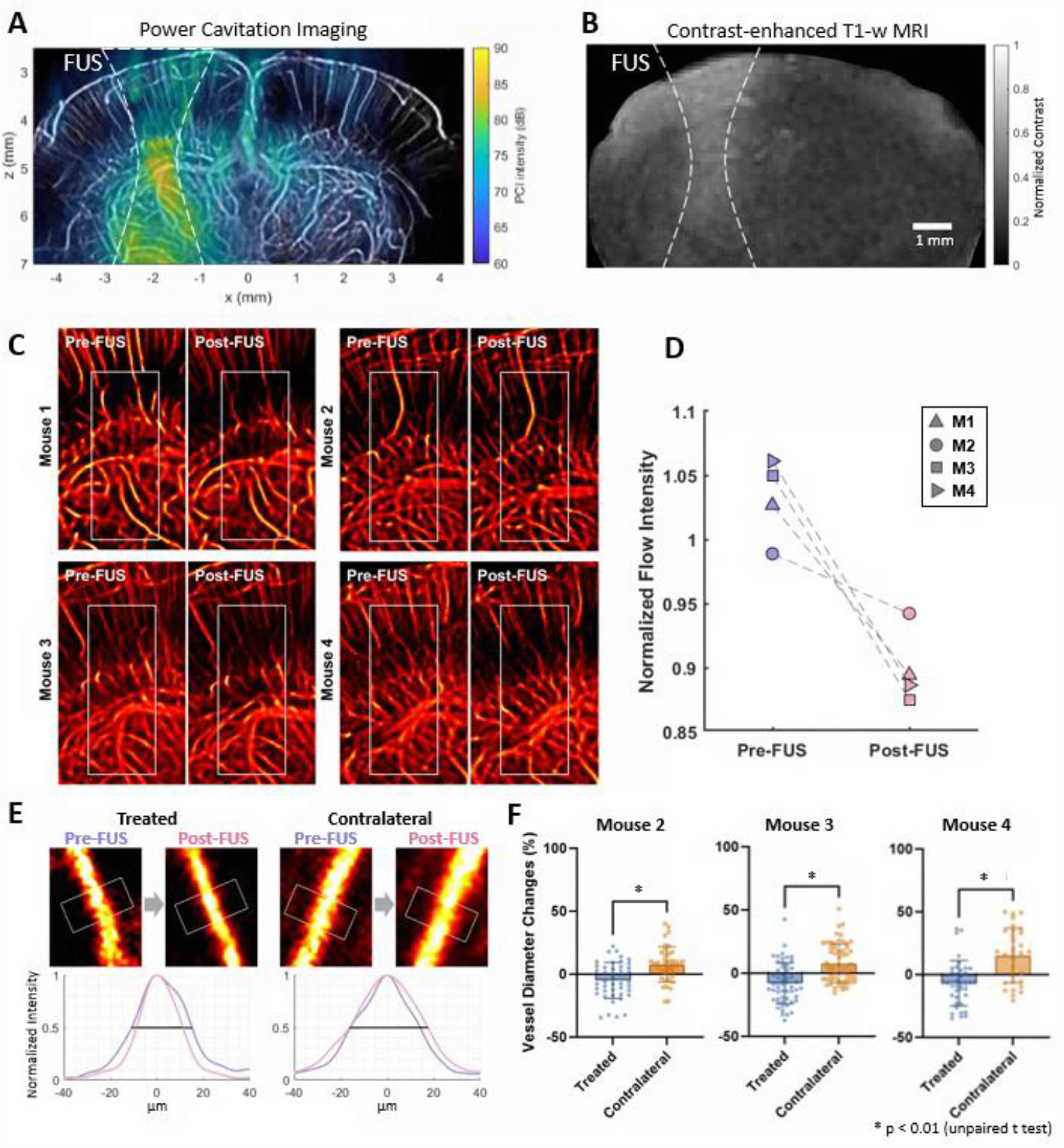
Cerebrovascular changes after FUS in the open-skull experiments. **A**) Power cavitation map obtained during FUS sonication. **B**) Resultant BBB opening verified in contrast-enhanced T1-weighted MRI. In A and B, the -6 dB contour of the synthesized pressure field of 5 foci is indicated by white dashed lines. **C**) Flow intensity maps before and after FUS at the treated region. **D**) Flow intensity at the treated region (white box in C) normalized by the contralateral region. Normalized flow intensity decreased following FUS. **E**) Representative vessel in the treated and contralateral regions for diameter measurements before and after FUS. Fifteen cross-sections were obtained within the segment (white box) and averaged to obtain a mean intensity profile. Its FWHM was measured as the diameter of the vessel. The full-width half-maximums of the mean intensity profiles of the pre-FUS (pink) and post-FUS (purple) were used for measuring the vessel diameter change. **F**) Vessel diameter changes after Mb-FUS in each mouse. Each data point represents the measurement from each vessel segment.

In ULM images, vessel segments were selected in both treated and contralateral regions, and the diameter of vessels before and after FUS was measured (Figures 3E). Although both vasoconstriction and vasodilation were found in both hemispheres, we found that vasoconstriction was more prevalent in the treated region, while vasodilation was more predominant in the contralateral region (Figure 3F).

### Transcranial detection of localized blood volume reduction

Transcranial ULM images before and after sonication with different pressures were presented in Figure 4A. A greater reduction in flow signal was observed in higher pressure cases. The BBBO size was linearly correlated with the cavitation dose obtained from PCD (Figure 4B). The average intensity measured within the white box in Figure 4A decreased as the pressure increased (Figure 4C), which was also visible in the ULM images.

**Figure 4:**
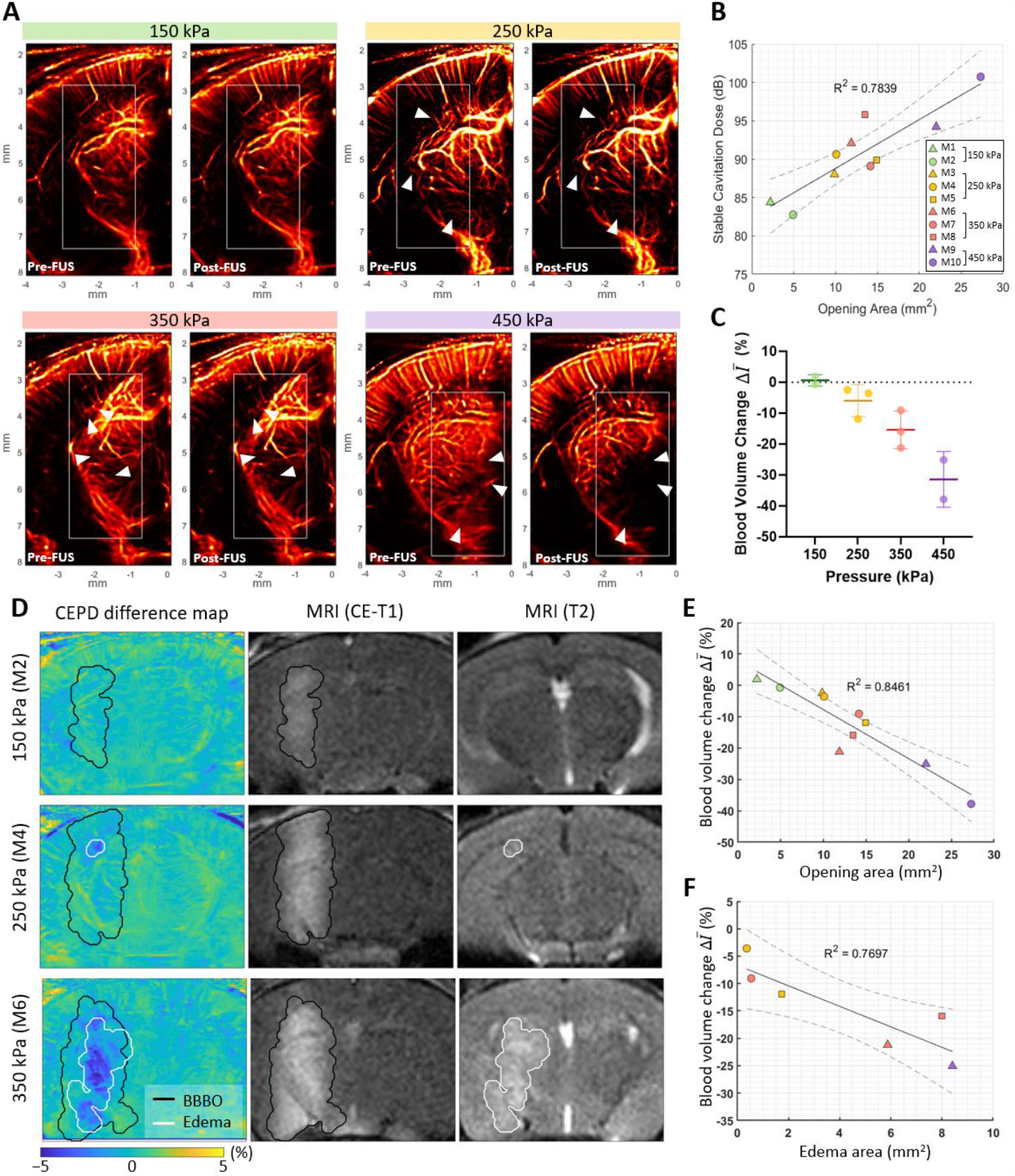
Blood volume reduction after Mb-FUS in the transcranial experiments. **A**) Representative pre-FUS (left) and post-FUS (right) ULM images for different acoustic FUS pressure groups (150, 250, 350, and 450 kPa). **B**) Stable cavitation dose detected by PCD with respect to the BBBO area. **C**) Blood volume change detected from ULM images for different pressure groups. **D**) Representative CEPD difference maps, CE-T1 MRI, and T2 MRI for pressure levels of 150, 250, and 350 kPa. BBBO region and edema region detected by CE-T1 and T2 MRI, respectively, are overlaid on the CEPD difference maps. **E**) Blood volume reduction with respect to the BBBO area. **F**) Blood volume reduction with respect to the edema area.

Power Doppler difference maps were obtained before and after BBBO and compared with CE-T1 and T2 MRI for different pressures (Figure 4D). BBBO and edema regions were quantified from MRIs, and contours were overlaid on the difference map. The blue-colored area in the difference map showed localized blood volume reduction (first column in Figure 4D). The maps showed again a greater reduction for the higher FUS pressure. Additionally, the localized reduced area in blue was matched with the edema region for the cases of 250 kPa and 350 kPa. However, for 150kPa, opening without edema case, the reduction was not noticeable, possibly due to the limited sensitivity of transcranial imaging.

Comparing the blood volume change with the size of BBBO and the size of edema, negative correlations were observed with the R^2^ of 0.85 and 0.77, respectively (Figure 4E and 4F).

## Conclusion

In this study, we established a new method for quantifying vascular changes after Mb-FUS using ultrasound flow imaging in mice. We found that Mb-FUS leads to a reduction in blood volume at the treated region, with vasoconstriction being more dominant than vasodilation. We also show that transcranial CEPD and ULM can detect localized blood reduction right after Mb-FUS. In addition, the CEPD difference map was spatially coincident with the edema region in T2-weighted MRI. These results showed the potential of ultrasound flow imaging as a monitoring tool of Mb-FUS to predict the BBBO size and to ensure the safety of the treatment.

## Reference

Bae, S., Liu, K., Pouliopoulos, A. N., Ji, R., & Konofagou, E. E. (2023). Real-time passive acoustic mapping with enhanced spatial resolution in neuronavigation-guided focused ultrasound for blood-brain barrier opening. IEEE Transactions on Biomedical Engineering, 70(10), 2874–2885.

Batts, A., Ji, R., Kline-Schoder, A., Noel, R., & Konofagou, E. (2022). Transcranial theranostic ultrasound for pre-planning and blood-brain barrier opening: a feasibility study using an imaging phased array in vitro and in vivo. IEEE Transactions on Biomedical Engineering, 69(4), 1481–1490. 10.1109/TBME.2021.3120919

Batts, A. J., Ji, R., Noel, R. L., Kline-Schoder, A. R., Bae, S., Kwon, N., & Konofagou, E. E. (2023). Using a novel rapid alternating steering angles pulse sequence to evaluate the impact of theranostic ultrasound-mediated ultra-short pulse length on blood-brain barrier opening volume and closure, cavitation mapping, drug delivery feasibility, and safety. Theranostics, 13(3), 1180–1197. 10.7150/thno.76199

Burgess, A., Nhan, T., Moffatt, C., Klibanov, A. L., & Hynynen, K. (2014). Analysis of focused ultrasound-induced blood-brain barrier permeability in a mouse model of Alzheimer’s disease using two-photon microscopy. Journal of Controlled Release, 192, 243–248. 10.1016/j.jconrel.2014.07.051

Burgess, M. T., Apostolakis, I., & Konofagou, E. E. (2018). Power cavitation-guided blood-brain barrier opening with focused ultrasound and microbubbles. Physics in Medicine and Biology, 63(6). 10.1088/1361-6560/aab05c

Chen, K.-T., Chai, W.-Y., Lin, C.-J., Lin, Y.-J., Chen, P., Tsai, H.-C., Huang, C.-Y., Kuo, J. S., Liu, H.-L., & Wei, K.-C. (2021). Neuronavigation-guided focused ultrasound for transcranial blood-brain barrier opening and immunostimulation in brain tumors. Science Advances, 7(6), eabd0772.

Cho, E. E., Drazic, J., Ganguly, M., Stefanovic, B., & Hynynen, K. (2011). Two-photon fluorescence microscopy study of cerebrovascular dynamics in ultrasound-induced blood-brain barrier opening. Journal of Cerebral Blood Flow and Metabolism, 31(9), 1852–1862. 10.1038/jcbfm.2011.59

Coviello, C., Kozick, R., Choi, J., Gyöngy, M., Jensen, C., Smith, P. P., & Coussios, C.-C. (2015). Passive acoustic mapping utilizing optimal beamforming in ultrasound therapy monitoring. The Journal of the Acoustical Society of America, 137(5), 2573–2585. 10.1121/1.4916694

Errico, C., Osmanski, B. F., Pezet, S., Couture, O., Lenkei, Z., & Tanter, M. (2016). Transcranial functional ultrasound imaging of the brain using microbubble-enhanced ultrasensitive Doppler. NeuroImage, 124, 752–761. 10.1016/j.neuroimage.2015.09.037

Errico, C., Pierre, J., Pezet, S., Desailly, Y., Lenkei, Z., Couture, O., & Tanter, M. (2015). Ultrafast ultrasound localization microscopy for deep super-resolution vascular imaging. Nature, 527(7579), 499–502. 10.1038/nature16066

Ji, R., Karakatsani, M. E., Burgess, M., Smith, M., Murillo, M. F., & Konofagou, E. E. (2021). Cavitation-modulated inflammatory response following focused ultrasound blood-brain barrier opening. Journal of Controlled Release, 337, 458–471. 10.1016/j.jconrel.2021.07.042

Jones, R. M., McMahon, D., & Hynynen, K. (2020). Ultrafast three-dimensional microbubble imaging in vivo predicts tissue damage volume distributions during nonthermal brain ablation. Theranostics, 10(16), 7211–7230. 10.7150/thno.47281

Kobus, T., Vykhodtseva, N., Pilatou, M., Zhang, Y., & McDannold, N. (2016). Safety Validation of Repeated Blood-Brain Barrier Disruption Using Focused Ultrasound. Ultrasound in Medicine and Biology, 42(2), 481–492. 10.1016/j.ultrasmedbio.2015.10.009

Konofagou, E. E. (2012). Optimization of the ultrasound-induced blood-brain barrier opening. Theranostics, 2(12), 1223–1237. 10.7150/thno.5576

Leinenga, G., & Götz, J. (2015). Scanning ultrasound removes amyloid-β and restores memory in an Alzheimer’s disease mouse model. Science Translational Medicine, 7(278), 278ra33. 10.1126/scitranslmed.aaa2512

Mainprize, T., Lipsman, N., Huang, Y., Meng, Y., Bethune, A., Ironside, S., Heyn, C., Alkins, R., Trudeau, M., Sahgal, A., Perry, J., & Hynynen, K. (2019). Blood-brain barrier opening in primary brain tumors with non-invasive MR-guided focused ultrasound: A clinical safety and feasibility study. Scientific Reports, 9(1), 1–7. 10.1038/s41598-018-36340-0

McMahon, D., O’Reilly, M. A., & Hynynen, K. (2021). Therapeutic agent delivery across the blood-brain barrier using focused ultrasound. Annual Review of Biomedical Engineering, 23, 89–113. 10.1146/annurev-bioeng-062117-121238

Pouliopoulos, A. N., Kwon, N., Jensen, G., Meaney, A., Niimi, Y., Burgess, M. T., Ji, R., McLuckie, A. J., Munoz, F. A., Kamimura, H. A. S., Teich, A. F., Ferrera, V. P., & Konofagou, E. E. (2021). Safety evaluation of a clinical focused ultrasound system for neuronavigation guided blood-brain barrier opening in non-human primates. Scientific Reports, 11(1), 1–32. 10.1038/s41598-021-94188-3

Raymond, S. B., Skoch, J., Hynynen, K., & Bacskai, B. J. (2007). Multiphoton imaging of ultrasound/Optison mediated cerebrovascular effects in vivo. Journal of Cerebral Blood Flow and Metabolism, 27(2), 393–403. 10.1038/sj.jcbfm.9600336

Roovers, S., Segers, T., Lajoinie, G., Deprez, J., Versluis, M., De Smedt, S. C., & Lentacker, I. (2019). The role of ultrasound-driven microbubble dynamics in drug delivery: From microbubble fundamentals to clinical translation. Langmuir, 35(31), 10173–10191. 10.1021/acs.langmuir.8b03779

Sheikov, N., McDannold, N., Sharma, S., & Hynynen, K. (2008). Effect of focused ultrasound applied with an ultrasound contrast agent on the tight junctional integrity of the brain microvascular endothelium. Ultrasound in Medicine and Biology, 34(7), 1093–1104. 10.1016/j.ultrasmedbio.2007.12.015

Willie, C. K., Colino, F. L., Bailey, D. M., Tzeng, Y. C., Binsted, G., Jones, L. W., Haykowsky, M. J., Bellapart, J., Ogoh, S., Smith, K. J., Smirl, J. D., Day, T. A., Lucas, S. J., Eller, L. K., & Ainslie, P. N. (2011). Utility of transcranial Doppler ultrasound for the integrative assessment of cerebrovascular function. In Journal of Neuroscience Methods (Vol. 196, Issue 2, pp. 221–237). Elsevier B.V. 10.1016/j.jneumeth.2011.01.011

Yang, Y., Zhang, X., Ye, D., Laforest, R., Williamson, J., Liu, Y., & Chen, H. (2019). Cavitation dose painting for focused ultrasound-induced blood-brain barrier disruption. Scientific Reports, 9, 2840. 10.1038/s41598-019-39090-9

